# Auditory Evoked Potentials Differ from Controls and Correlate with Aspects of Cognition in Kabuki Syndrome and Rubinstein-Taybi Syndrome

**DOI:** 10.64898/2026.11.04.764287

**Authors:** A Kalinousky, E Bucklin, R Ng, L Meier, H Jiddou, P Fanara, JB Ewen, JR Harris

## Abstract

**Objective:** Kabuki syndrome (KS) and Rubinstein-Taybi syndrome (RSTS) are two disorders caused by dysfunctional components of the epigenetic machinery and both result in intellectual disability. Mouse models of both KS and RSTS have seen reversal of cognitive impairment with epigenetic treatments. However, a great need exists for biomarkers in these disorders to improve the likelihood of successful clinical trials. This study aimed to investigate cortical auditory evoked potentials (AEPs) in KS and RSTS so that it could potentially be developed into a response biomarker in the future.

**Methods:** The amplitude of the N1-P2-N2 complex was measured for passive listening to monotonal auditory stimuli in individuals with KS, with RSTS, and typically developing controls. A standardized cognitive assessment battery was performed and specific subtests were correlated with AEP values.

**Results:** Groups differed significantly in P2, N2, and P2-N2 amplitude although RSTS was consistently more different from typically developing controls than was KS. The mid- to late-phase AEP components (P2 and N2) also correlate with cognition within and across these groups particularly with language whereas earlier components N1 and P2 correlate more with measures of executive functioning.

**Conclusion:** Late-phase components of AEPs differ in KS and RSTS from each other and from controls. KS and RSTS differ from controls in P2 and N2 amplitude in the same direction and both show a positive correlation between language and P2 amplitude and a negative correlation between language and N2 amplitude. Together they also show an association with N1 and P2 amplitude and executive functioning.

**Significance:** AEPs may turn out to be a valuable response biomarker in KS and RSTS individually and together.

## 1. Introduction

### 1.1 Mendelian Disorders of the Epigenetic Machinery: Kabuki Syndrome and Rubinstein-Taybi Syndrome

Kabuki syndrome (KS) and Rubinstein-Taybi syndrome (RSTS) are both examples of Mendelian disorders of the epigenetic machinery (MDEMs) (Fahrner and Bjornsson, 2019). MDEMs are a group of rare disorders that almost universally result in neurodevelopmental disorders (Ng, R. et al., 2023) KS and RSTS both have mouse models and previously published work has shown that the cognitive impairment in the KS and RSTS mouse models can be corrected with epigenetic modification therapies (Bjornsson et al., 2014b; Benjamin et al., 2017; Bjornsson et al., 2014a; Alarcón et al., 2004). Despite this, neither disorder has had a large-scale clinical trial in humans and one reason is the lack of established biomarkers in these conditions.

Kabuki syndrome (KS) is a MDEM occurring in about 1 in every 32,000 births characterized by distinctive facial gestalt, postnatal growth deficiency, hypotonia, and multiple congenital anomalies (Adam et al., 1993) The majority of KS is caused by heterozygous pathogenic variants in *KMT2D* (Ng, S. B. et al., 2010). KS resulting from *KMT2D* variants is known as Kabuki syndrome type 1 (KS1; MIM 147920). Mutations in *KDM6A* are the less common cause of KS known as Kabuki syndrome type 2 (KS2; MIM 300867) (Banka et al., 2014) Mutations in *KMT2D* or *KDM6A* both result in heterochromatin and similar widespread changes in gene expression and KS1 and KS2 have overlapping phenotypes. In both forms of KS, the neurodevelopmental phenotype is the most prevalent and the most impairing (Theodore-Oklota et al., 2021) The neurocognitive and neurobehavioral phenotype of Kabuki syndrome has been previously described in the literature – mild intellectual disability with disproportionate and significant weaknesses in visuospatial construction, perception and memory(Harris et al., 2019; Dongen et al., 2019) and prosocial behavior with features of attention-deficit hyperactivity disorder especially early in life and significant anxiety that worsens with age (Kalinousky, A. J. et al., 2024; Kalinousky, Allison J. et al., 2022)

Rubinstein-Taybi syndrome (RSTS) is characterized by short stature, cognitive impairment, specific facial features, growth impairment, and broad thumbs and great toes (Stevens, 1993). Like KS, RSTS is another example of an MDEM as mutations in 2 different genes that creates convergent epigenetic dysfunction and results in the same disorder. RSTS1 (MIM 180849) is caused by heterozygous pathogenic variants in *CREBBP* whereas RSTS2 (MIM 613684) is caused by heterozygous pathogenic variants in *EP300* (Stevens, 1993). CREBBP and EP300 are closely related histone acetyltransferases which also alter gene expression. RSTS1 and RSTS2 have nearly identical phenotypes although RSTS2 typically has milder facial features and less severe intellectual disability (Fergelot et al., 2016) The cognitive and neurobehavioral phenotype of RSTS has similarities to KS – intellectual disability and poor visuospatial skills with strong social drive but significant ADHD and anxiety that is more apparent with age (Lacombe et al., 2024; Qu’d et al., 2023). RSTS typically has more severe intellectual disability and weaker language skills than KS and more significant behavioral dysregulation (Lacombe et al., 2024; Qu’d et al., 2023)

Mouse models of KS1 and RSTS1 demonstrate hippocampal memory deficits which have been ameliorated in postnatal life with epigenetic therapies (Bjornsson et al., 2014b; Benjamin et al., 2017; Bjornsson et al., 2014a; Alarcón et al., 2004). However before robust clinical trials can be undertaken, there is a great need for biomarkers that reflect the underlying cognitive processes and differences in KS and RSTS.

### 1.2 Auditory Evoked Potentials

Auditory evoked potentials (AEPs) are temporally precise measurements of neurophysiologic reactions to auditory stimuli that are captured using EEG. AEPs provide an quantitative and non-invasive measure of neurologic processing that can be passive without complicated instructions or active attention to tasks. This makes AEP a very good tool for studying populations with intellectual disability. Many studies have shown differences in neurodevelopmental groups and typically developing individuals in AEPs (Ethridge et al., 2019; Proteau-Lemieux et al., 2025; Davies et al., 2010; Saby et al., 2023; Schwartz et al., 2023). Some genetic intellectual disability syndromes such as Rett syndrome and Fragile X syndrome are using AEPs as a validated biomarker in clinical trials (Ethridge et al., 2019; Saby et al., 2021). To date, no studies have examined evoked potentials in KS or RSTS populations.

The present study seeks to determine the potential utility of evoked potentials (EPs) as a clinically relevant biomarker of cortical function in KS and RSTS.

## 2. Methods

### 2.1 Participants

Participants with KS and RSTS were recruited through clinics at Kennedy Krieger Institute and Johns Hopkins Medical Institute, by contacting previous study participants who consented to be contacted, through attendance at syndrome-based conferences, through partnerships with Kabuki Syndrome Foundation and Rubinstein-Taybi Syndrome Children’s Foundation, and advertisement on support group websites. Inclusion criteria for participants was documentation of a pathogenic or likely pathogenic variant in *KMT2D, KDM6A, CREBBP*, or *EP300* combined with clinical diagnosis of the respective disorder. Exclusion criteria is presence of another genetic syndrome that affects neurodevelopment, hearing loss that is in the severe range even with correction, and inability/unwillingness to participate in EEG. Typically developing (TD) control participants were recruited from previous control populations who consented to be contacted about further research, from other clinics at the authors’ institutions, and many were family members of individuals with KS and RSTS. Exclusion criteria for controls included neurodevelopmental disability, history of brain malformation or moderate to severe traumatic brain injury, hearing loss that is in the severe range even with correction, inability/unwillingness to participate in EEG. and presence of any genetic syndrome.

### 2.2 Cognitive and Behavioral Measures

All participants underwent cognitive and behavioral assessment on the same day as the AEP. Scores were generated based on this testing for comparison with AEP parameters. All participants had domain scores calculated for Language, Nonverbal IQ, Visual Motor Integration, Visual Perception, Attention and Inhibition, Cognitive Flexibility, Behavioral Issues, and Adaptive Functioning. Language domain scores were calculated using either the Verbal Comprehension Index of the age-appropriate Wechsler evaluation (Wechsler, 2003) or from the Peabody Picture Vocabulary Test Standard Score. Nonverbal IQ was calculated using either the age-appropriate Wechsler tests or the Leiter International Performance Scale -3. Visual Motor Integration and Visual Perception scores were standard scores from the Beery-Buktenika Test of Visual Motor Integration(Beery and Buktenica, 2004). For Attention and Inhibition and Cognitive Flexibility we used the age-corrected standard scores from the NIH Toolbox Flanker Inhibition Test and the NIH Toolbox Dimensional Change Card Sort Test respectively (Wechsler, 2003; Shields et al., 2020). Behavioral Issues used the Child Behavior Checklist or Adult Behavior Checklist Total Problems Score (Achenbach T.M., 2000). For the Adaptive Functioning domain, the General Adaptive Composite from the Adaptive Behavior Assessment Scale was used.

### 2.3 Auditory Evoked Potentials

EEG data were recorded using a 47-channel WaveGuard cap system and asa-lab amplifier. To capture the AEP data, The stimuli were presented using E-Prime 2.0 (Psychology Software Tools, Inc.) connected to a chronos box connected to a stimtracker and then connected to speakers and the amp. Participants were seated in a soundproof booth with binaural speakers. Auditory stimuli were identical 1000 Hz tones presented at 70 dB and with a jittered interstimulus interval. A caregiver and/or child life specialist can be in the room with the participant in order to encourage participants not to speak and to remain through the whole protocol. Participants can watch a silent movie or video, look at pictures, or play with a silent toy. Raw data will be recorded continuously.

EEG data was preprocessed using HAPPE, an automated preprocessing program.(Gabard-Durnam et al., 2018) This serves to clean the data by removing known artifacts and filtering noise. It also separates the data into individual evoked potential events. Data is filtered using a 30 Hz low-pass filter, segmented on stimulus onset to include a 100 ms prestimulus baseline and a 600 ms post-stimulus interval, and screened for ocular and movement artifacts.

The six electrodes in the electrode group surrounding Cz were all averaged together to get the multi-average values for N1, N2, and P2 as has been previously published.(Zhang and Kappenman, 2024) The N1 was identified between 70 and 170 ms and the P1 was scored between 20 and 80 ms. The peak-to-peak amplitude of the N1 component was defined as the difference in μV between the N1 peak amplitude and the P1 peak amplitude. The P2 component was identified as the most positive peak between 130 and 270 ms after the stimulus onset and peak-to-peak amplitude was defined as the difference in μV between the N1 peak and the P2 peak. The N2 component was identified as the most negative peak between 200 and 375 ms after the stimulus onset and the peak-to-peak amplitude was defined as the difference in amplitude between the P2 peak and the N2 peak.

### 2.4 Statistical Analysis

All statistical analyses were conducted using R for Windows (R Foundation for Statistical Computing, Vienna, Austria). Group differences in AEP parameters were analyzed using ANCOVA with age and sex as covariates for each AEP measure. The Benjamini–Hochberg correction was applied across the AEP outcomes. For the AEP-cognitive correlations, we used separate linear regressions to predict a cognitive score from one AEP measure, adjusting for age and sex. In the pooled KS+RSTS models, it also adjusted for diagnosis.

## 3. Results

### 3.1 Participants

35 participants with KS, 23 participants with RSTS and 40 TD participants were recruited for this study. Figure 1 shows the overall participant flow. For KS, 4 participants did not behaviorally comply with capping. Of the 31 participants who were capped, 26 had AEPs used in analysis. Of the 5 excluded, three were excluded for excessive artifact and/or lack of a valid AEP (defined as a valid N1 wave occurring between 80-130 ms from stimulus) and two were excluded because they had a present AEP but not at Cz. For RSTS, another 4 participants did not behaviorally comply with capping. Of the 19 participants who were capped, 11 had AEPs used in analysis. Of the 8 excluded, two were excluded for excessive artifact and/or lack of a valid AEP and 7 were excluded because they had a present AEP but not at Cz. All 40 TD participants were able to be capped and 34 had AEPs used in analysis. Of the 6 excluded, two were excluded for excessive artifact and/or lack of a valid AEP and 4 were excluded because they had a present AEP but not at Cz. The demographics of the participants analyzed is shown in Table 1.

**Table 1:** Participant Demographics.

| Group | Age in Months M (SD) | % Female | NVIQ | Language | VMI | VP | CBCL | BRIEF | ABAS | NIHTB Flanker | NIHTB Card Sort |
|---|---|---|---|---|---|---|---|---|---|---|---|
| KS (n=26) | 223 (100) | 69 | 71 | 79 | 58 | 60 | 62 | 67 | 72 | 61 | 76 |
| RSTS (n=11) | 142 (42) | 27 | 66 | 64 | 53 | 66 | 67 | 75 | 67 | 62 | 63 |
| TD (n=34) | 209 (111) | 56 |  |  |  |  |  |  |  |  |  |

KS = Kabuki syndrome, RSTS = Rubinstein-Taybi syndrome, TD = typically developing, NVIQ = Nonverbal IQ standard score from age-appropriate Wechsler test or Leiter, Language = standard score from age-appropriate Wechsler test or Peabody Picture Vocabulary Test, VMI = Beery-Buktenika Visual Motor Integration standard score, VP = Beery-Buktenika Visual Perception standard score, CBCL = Child Behavior Checklist (could also be Adult Behavior Checklist) Total Problems Score, BRIEF = Behavior Rating Inventory of Executive Functioning Global Executive Composite Score, ABAS = Adaptive Behavior Assessment Scales General Adaptive Composite Score, NIHTB Flanker = NIH Toolbox Flanker Inhibition test age-corrected standard score, NIHTB Card Sort = NIH Toolbox Dimensional Change Card Sort test age-corrected standard score

**Figure 1:**
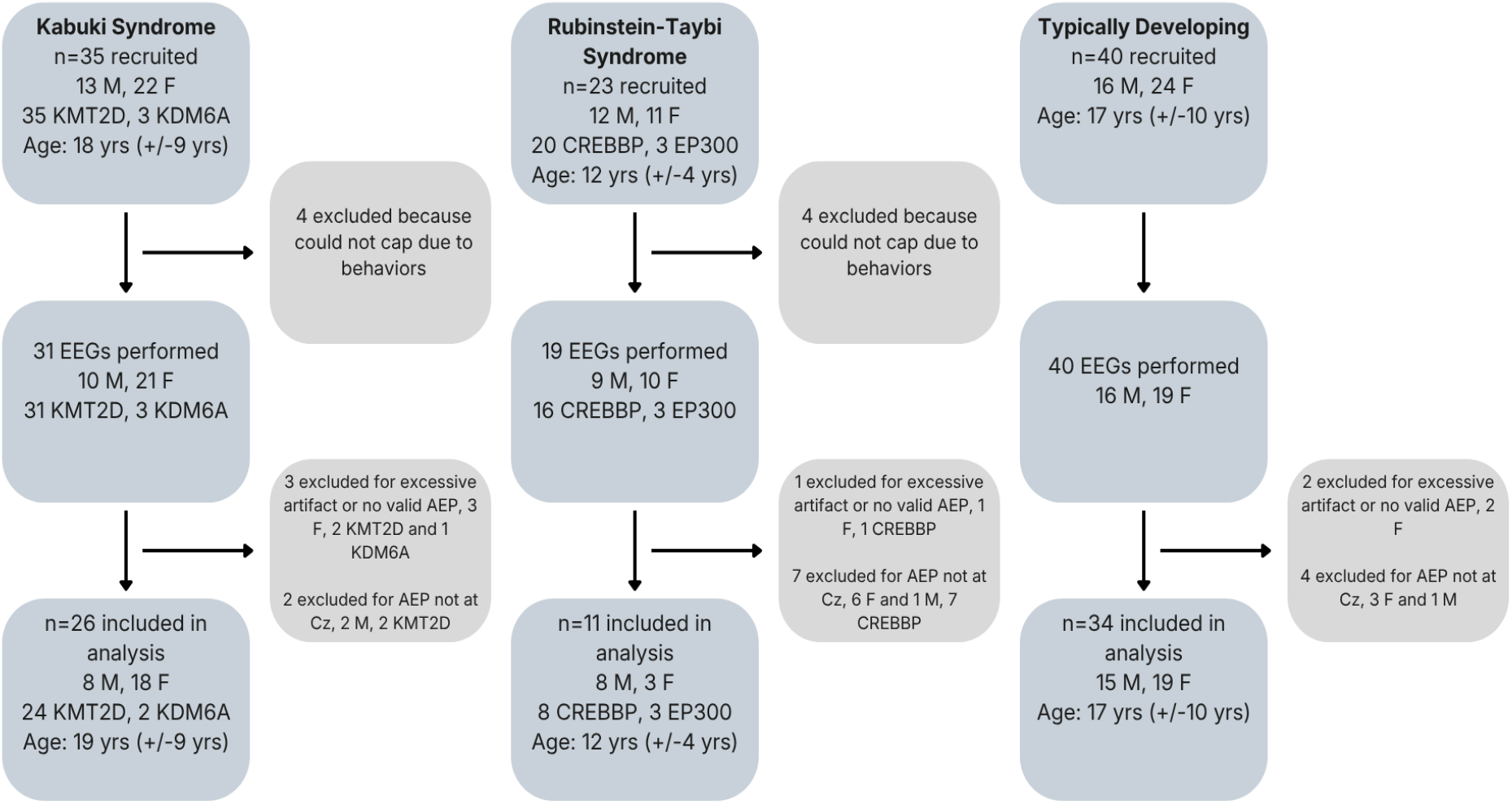
Participant Floechart. Demographic and clinical information for all participants enrolled in the study, reasons for exclusion, and final samples for the AEP analyses.

There were no statistically significant differences sex between groups but the RSTS group was significantly younger than KS or TD. Individual genotypes for KS and RSTS participants are shown in Supplementary Table 1. In summary, for KS, 24 analyzed participants had *KMT2D* variants (14 frameshift or nonsense variants and 10 missense or splice site variants) and 2 had *KDM6A* variants. For RSTS, 8 analyzed participants had *CREBBP* variants (14 frameshift or nonsense variants and 10 missense or splice site variants) and 3 had *EP300* variants. Of note, all of those excluded from the RSTS analysis for lack of AEP at Cz had *CREBBP* variants. The overall diagnosis groups did not vary significantly between age or sex distribution.

### 3.2 AEP Analysis

The grand average waveform for all three groups is shown in Figure 2. There were no significant differences seen in males vs females within or across diagnoses. Age was controlled in our models but interestingly, age was a significant factor in P2 and N2 amplitude for TD but not for KS and RSTS. Amplitude was significantly more negative in N2 for both KS and RSTS vs TD and also for KS+RSTS pooled data vs TD. The absolute value of the P2 to N2 difference was also significantly larger for RSTS, and KS+RSTS versus controls (Fig 2 and Table 2). There were no significant group differences in any other AEP amplitudes. Given the small sample sizes we did not look at KS1 vs KS2 or RSTS1 vs RSTS2.

**Table 2:** AEP Amplitude Parameters Compared Between KS, RSTS, and TD.

| AEP Parameter | Comparison | Estimate | Conf Lower | Conf Upper | p-value | p_adjusted |
|---|---|---|---|---|---|---|
| N1 | KS vs TD | 4.82E-07 | -1.53E-07 | 1.12E-06 | 0.13 | 0.24 |
|  | RSTS vs TD | 1.42E-07 | -8.26E-07 | 1.11E-06 | 0.77 | 0.77 |
|  | KS+RSTS vs TD | 3.88E-07 | -2.17E-07 | 9.93E-07 | 0.2 | 0.34 |
| N1-P2 | KS vs TD | -2.77E-07 | -1.58E-06 | 1.03E-06 | 0.67 | 0.67 |
|  | RSTS vs TD | 5.37E-07 | -1.44E-06 | 2.51E-06 | 0.59 | 0.73 |
|  | KS+RSTS vs TD | -2.42E-08 | -1.33E-06 | 1.29E-06 | 0.97 | 0.97 |
| P2 | KS vs TD | 2.05E-07 | -6.57E-07 | 1.07E-06 | 0.64 | 0.67 |
|  | RSTS vs TD | 6.79E-07 | -6.25E-07 | 1.98E-06 | 0.3 | 0.5 |
|  | KS+RSTS vs TD | 3.64E-07 | -4.96E-07 | 1.22E-06 | 0.4 | 0.5 |
| P2-N2 | KS vs TD | 8.37E-07 | -3.02E-07 | 1.97E-06 | 0.15 | 0.24 |
|  | RSTS vs TD | 3.13E-06 | 1.43E-06 | 4.82E-06 | 0.0006* | 0.0015* |
|  | KS+RSTS vs TD | 1.56E-06 | 3.88E-07 | 2.74E-06 | 0.0099* | 0.025* |
| N2 | KS vs TD | -6.31E-07 | -1.09E-06 | -1.69E-07 | 0.0084* | 0.042* |
|  | RSTS vs TD | -2.45E-06 | -3.22E-06 | -1.68E-06 | 1.12e-07* | 5.62e-07* |
|  | KS+RSTS vs TD | -1.20E-06 | -1.74E-06 | -6.60E-07 | 3.61e-05* | 0.00018* |

AEP Estimates between groups. Direction is first comparator vs second. N1-P2 and P2-N2 are absolute value magnitudes.

**Figure 2:**
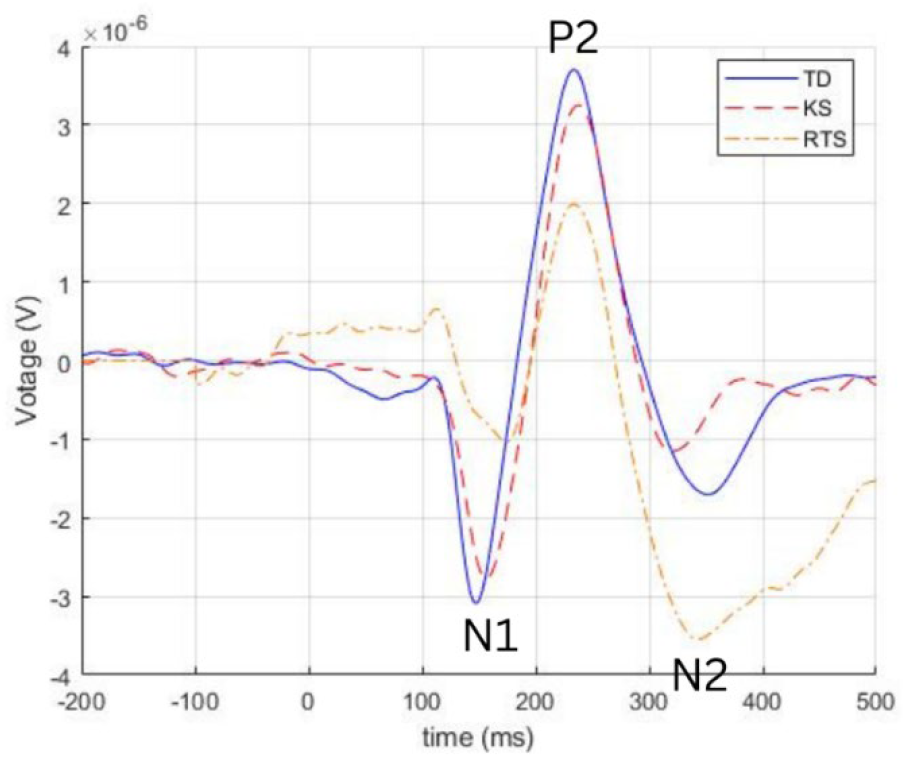
Grand Average Waveform. Grand average waveform for each group at Cz. N1, P2, and N2 components are labeled.

**Figure 3:**
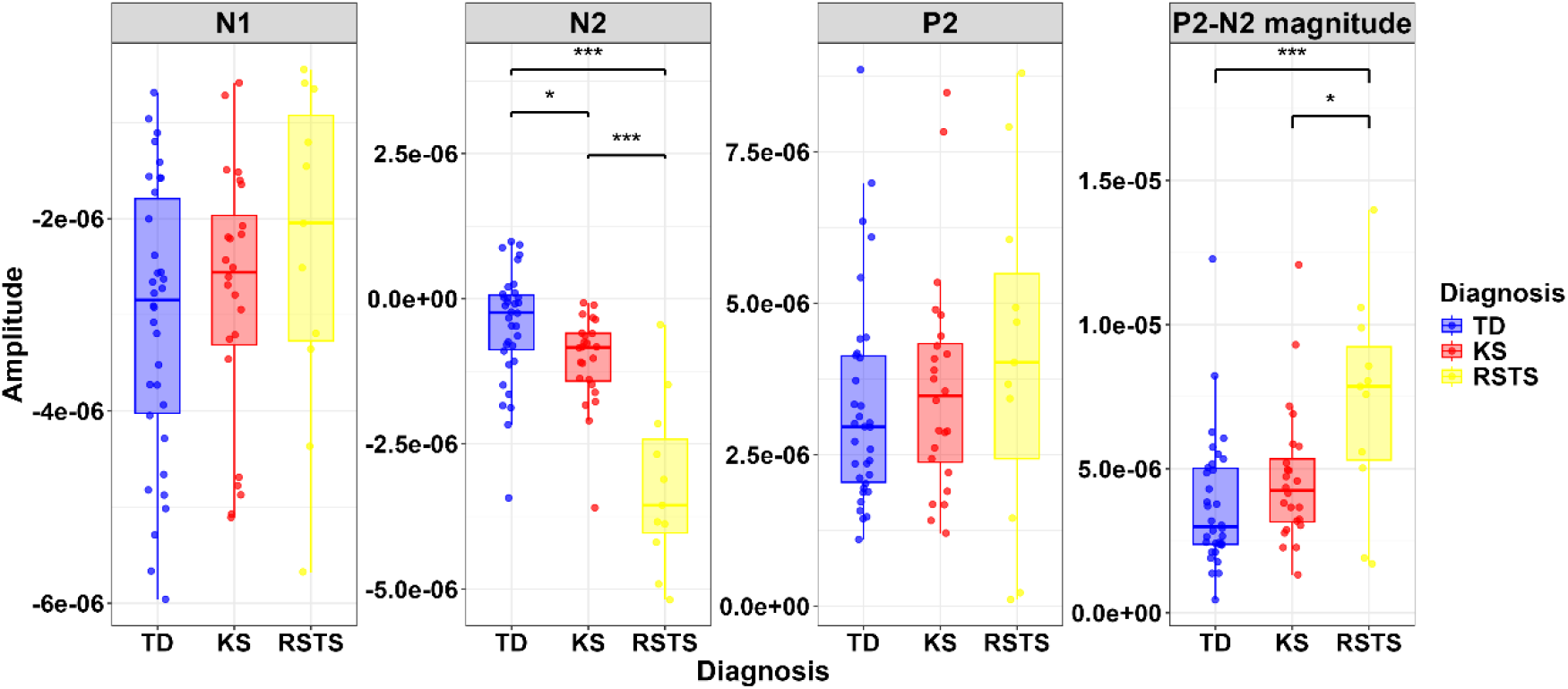
AEP Parameter Amplitudes KS, RSTS, and TD. Bar graphs showing the median values and inter-quartile range for AEP amplitudes for KS, RSTS, and TD. The amplitude of the N2 and P2-N2 magnitude significantly differ in the RSTS group compared to KS and TD. The amplitude of N2 is significantly lower in KS compared to TD.

When looking across all cognitive domains, 4 had correlations with the AEP parameters – Language, VMI, BRIEF and NIHTB Flanker Inhibition. For the KS group, P2, P2-N2, and N2 significantly correlated with language in this sample. P2 had a positive association and N2 had a negative association. The absolute value of P2-N2 positively correlated with language. N2 also had a negative correlation with VMI. For the RSTS group, nothing achieved significance with only 11 participants. However NIHTB Flanker trended toward a significant association with N1, N1-P2, and P2. When looking at the KS and RSTS group together, NIHTB Flanker was significantly correlated with N1, N1-P2, P2 and P2-N2. Flanker had a negative correlation with N1 and a positive correlation with P2 and with the magnitudes of N1-P2 and P2-N2. BRIEF was significantly correlated with N1-P2, P2, and P2-N2. Language was significantly correlated with N1-P2, P2, P2-N2, and N2. It was positively correlated with P2 and the magnitudes of N1-P2 and P2-N2 and negatively with N2. VMI showed a significant negative correlation with N2.

Given that this study was designed to explore AEP associations in KS and RSTS and not to definitively validate AEPs as a biomarker, and given the small sample sizes we did not correct for multiplicity in these correlation analyses. The associations for the most correlated cognitive parameters – Language, VMI, BRIEF, and NIHTB Flanker – are shown in Figure 4. Every association is shown in Supplementary Table 2.

**Figure 4:**
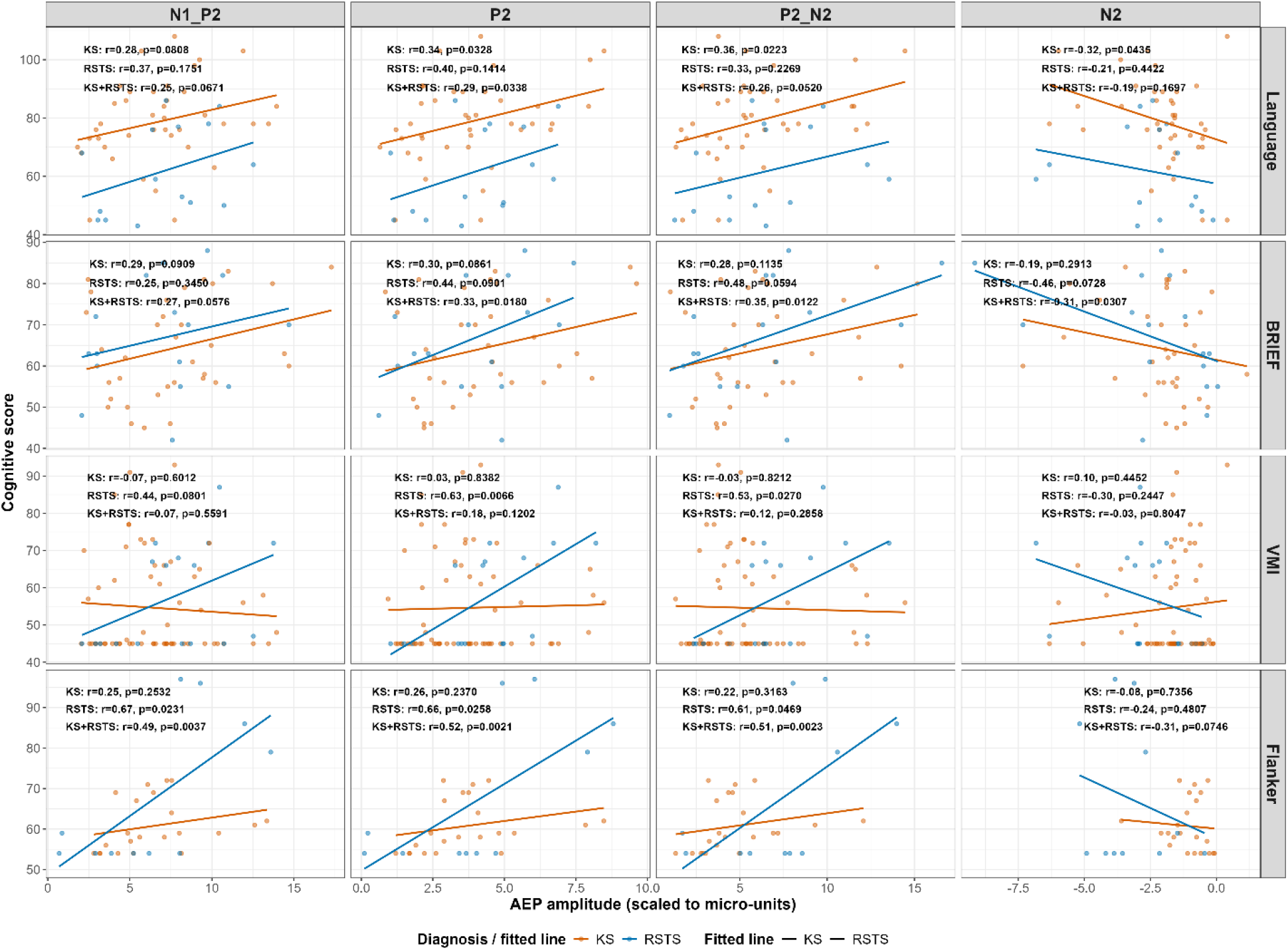
Cognitive Association with AEP Parameters Within and Across Groups. Correlations of 4 AEP parameters (N1 excluded because of lack of any trends) and 4 cognitive domains that most contribute to prediction for KS alone, RSTS alone and KS+RSTS. Language = standard score from age-appropriate Wechsler test or Peabody Picture Vocabulary Test, VMI = Beery-Buktenika Visual Motor Integration standard score, BRIEF = Behavior Rating Inventory of Executive Functioning Global Executive Composite Score, NIHTB Flanker = NIH Toolbox Flanker Inhibition test age-corrected standard score

## 4. Discussion

This is the first study to examine AEP differences in two related but distinct MDEMs – KS and RSTS. We used a passive paradigm to ensure that it could be applicable to all individuals in these two syndromes regardless of degree of cognitive impairment. Other genetic neurodevelopmental disorders have shown syndrome-specific differences in AEPs. For example, Fragile X show larger magnitude N1 and P2 amplitudes thought to be associated with excessive cortical excitability (Ethridge et al., 2019) In contrast, AEPs in Rett syndrome show dampened magnitudes of P1, N1, and P2 thought to be indicative of decreased healthy neurologic functioning (Saby et al., 2021)

Typically in AEPs, N1 is thought to reflect early cortical detection and sensory processing of the auditory stimulus. P2 is associated with perceptual analysis of the sound and the formation or refinement of auditory representations. N2 is commonly linked to higher-order cognitive processes such as stimulus discrimination, novelty detection, conflict monitoring, or attention-related processing. Therefore, the later AEP parameters – P2 and N2 – are thought to be more related to cognition than N1 which is more just sensory detection. Here we provide good evidence of increased magnitude of P2 and N2 in KS and RSTS compared to typically developing controls with the magnitude of RSTS greater than KS in both. The P2-N2 magnitude was also, of course, increased. To further support that P2 and N2 are the important parameters in these conditions, our data shows that P2 and N2 are significantly correlated with aspects of cognition much more so than other AEP parameters. Interestingly, higher language and visual motor integration scores were associated with larger magnitude P2, N2, and P2-N2. This is particularly true for KS which had the more robust sample size. The association with language and P2, N2 and P2-N2 holds true when KS and RSTS are looked at together as a single group.

For RSTS, the greatest signal is a positive correlation with P2 amplitude and the NIH Toolbox Flanker Inhibition which also holds true when the KS and RSTS groups are combined. One possible explanation for the fact that the magnitude of P2 and N2 is larger in KS and RSTS than controls but also positively correlated with better cognition is that individuals with KS and RSTS have inefficient executive functioning and sensory and cognitive processing and therefore require recruitment of additional neural resources to complete the processing and regulate attention when compared to controls. The better their brains can achieve this additional network recruitment, the better their cognition. This hypothesis is particularly interesting given the large variability in topology of AEPs that required so many RSTS AEPs to be excluded from the analysis in this study. This could constitute evidence that differing neural recruitment is taking place. This would need testing and confirmation in larger studies.

An aspect of attention and executive functioning – NIHTB Flanker Inhibition – was correlated with the earlier AEP parameters N1 and P2. This could indicate a pattern where the executive function and degree of ADHD symptoms of individuals with KS and RSTS affect the earlier AEP parameters associated more with sensory processing and more cognitive domains like language and visuomotor integration correlate with the later parameters.

There are a number of limitations to this study. First, we had an unusually high number of RSTS participants whose data was not able to be analyzed. While we expect more behavioral failures in RSTS given their higher rate of behavioral challenges, the number of participants with an AEP present somewhere other than Cz was a surprise. This warrants further investigation into the topology of AEPs in RSTS especially in light of the potential differing neural recruitment represented by the P2 and N2 amplitude findings in KS and RSTS. Second, while all participants have prospectively collected cognitive data in specific domains, some participants were tested using different measures as part of different research protocols and with small sample sizes this could affect analysis especially looking across groups.

Future directions include better characterizing the morphology and topology of AEPs in both KS and RSTS. This is an initial study in these conditions and needs to be reproduced. Topology differences particularly for RSTS should be investigated as mentioned above. Additionally, even if the findings of consistent differences in P2 and N2 and correlation with severity are further crystallized and replicated, several steps are still required for establishment of AEPs as a biomarker for KS and RSTS. Next step studies should have larger sample sizes in all genotypes with a more consistent cognitive battery. Showing consistency across sites would also be important. After that, AEPs would need to be shown to be responsive to intervention and the way AEPs change over time in individuals with KS and RSTS will need to be characterized., and across different sites would help validate AEPs as a potential biomarker in these conditions.

## 5. Conclusions

This study lays the groundwork for establishing differences between KS and RSTS and controls and early evidence for cognitive correlation with the P2-N2 complex signaling as particularly important in these conditions.

