## Supplemental Table 1 for "Auditory Evoked Potentials Differ from Controls and Correlate with Aspects of Cognition in Kabuki Syndrome and Rubinstein-Taybi Syndrome"

Supplementary Table 1 Genetic variants of participants included in AEP analysis

| ID | Sex | Age | Gene | c. | p. |
| --- | --- | --- | --- | --- | --- |
| KS001 | M | 15 | KMT2D | c.3190dupG |  |
| KS002 | M | 28 | KMT2D | c.14006C>G | p.Ser4669X |
| KS003 | F | 36 | KMT2D | c.14580dupT |  |
| KS005 | F | 27 | KMT2D | c.15536G>A | p.R5179H |
| KS006 | F | 18 | KMT2D | c.4265 G>A | p.W1422X |
| KS007 | F | 15 | KMT2D | IVS50+5G>A |  |
| KS008 | F | 12 | KMT2D | c.5278_5279delAA | K1760EfsX27 |
| KS009 | M | 20 | KMT2D | c.13432dupC | p.R4478fs |
| KS010 | F | 30 | KMT2D | c. 15536G>A | p. R517911 |
| KS011 | F | 20 | KMT2D | c. 7481_7482insT |  |
| KS012 | M | 22 | KMT2D | c.10813C>T | p. Q3605X |
| KS013 | F | 16 | KMT2D | c.7643dupA |  |
| KS014 | F | 13 | KMT2D | c.15251T>C | p.Leu5084Pro |
| KS016 | F | 12 | KMT2D | c.16018C>T | p.Arg5340* |
| KS017 | M | 20 | KMT2D | c.15142C>T | p.R5048C |
| KS019 | F | 20 | KMT2D | c.15626G>T | p. Gly5209Val |
| KS021 | M | 21 | KMT2D | c.15143G>A | R5048H |
| KS022 | F | 35 | KMT2D | c.14641C>T | p.Q4881* |
| KS023 | F | 25 | KMT2D | c.13681del | p.L4561Cfs*4 |
| KS024 | M | 12 | KMT2D | c.15535C>T | p.R5179C |
| KS027 | F | 13 | KDM6A | c.3978_3981delACCA | p.P1327LfsX28 |
| KS028 | F | 5 | KMT2D | c.4616_4617dupCA | p.E1540QfsX23 |
| KS029 | F | 14 | KMT2D | c.8366+2T>A |  |
| KS038 | M | 26 | KMT2D | c.5124_5125delAC |  |
| KS040 | F | 11 | KMT2D | c.15536G>A | p.Arg5179His |
| KS047 | M | 11 | KMT2D | c.2533delC | p.Arg845Glyfs*85 |
| RTS001 | F | 9 | CREBBP | c.3874C>T |  |
| RTS002 | F | 11 | CREBBP | c.5050T>C |  |
| RTS003 | M | 19 | EP300 | c.3070_3074delAAAGA |  |
| RTS005 | M | 12 | CREBBP | c.1823+5G>A |  |
| RTS006 | F | 6 | CREBBP | c.1823+5G>A |  |
| RTS009 | F | 18 | EP300 | c.1517delT | p.M506RfsX4 |
| RTS010 | M | 12 | CREBBP | c.5902C>T |  |
| RTS011 | M | 11 | CREBBP | c.6404delinsTGC | p.Gln2135Leufs*9 |
| RTS017 | M | 14 | CREBBP | deletion of exon 1 |  |
| RTS019 | M | 9 | CREBBP | c.4394+4A>G |  |
| RTS022 | M | 9 | EP300 | c.4783T>G |  |
