## Supplemental Table 2 for "Auditory Evoked Potentials Differ from Controls and Correlate with Aspects of Cognition in Kabuki Syndrome and Rubinstein-Taybi Syndrome"

| Supplementary Table 2: AEP Parameter Correlations with Cognition | | | | |
| --- | --- | --- | --- | --- |
| Group | AEP Parameter | Cognitive Domain | Estimate | Raw p-value |
| KS | N1 | NVIQ | -0.95 | 0.62 |
|  |  | Language | -0.65 | 0.57 |
|  |  | VMI | 1.39 | 0.30 |
|  |  | VP | -0.38 | 0.69 |
|  |  | CBCL | -2.52 | 0.09 |
|  |  | BRIEF | -1.27 | 0.22 |
|  |  | ABAS | -0.57 | 0.69 |
|  |  | NIHTB Flanker | -0.15 | 0.23 |
|  |  | NIHTB Card Sort | 2.22 | 0.16 |
|  | N1-P2 | NVIQ | 0.59 | 0.47 |
|  |  | Language | 1.22 | 0.09 |
|  |  | VMI | -0.8 | 0.31 |
|  |  | VP | -0.53 | 0.65 |
|  |  | CBCL | 1.07 | 0.12 |
|  |  | BRIEF | 0.85 | 0.14 |
|  |  | ABAS | 0.42 | 0.60 |
|  |  | NIHTB Flanker | -0.45 | 0.45 |
|  |  | NIHTB Card Sort | -1.31 | 0.14 |
|  | P2 | NVIQ | 1.24 | 0.38 |
|  |  | Language | 2.74 | 0.01* |
|  |  | VMI | -1 | 0.36 |
|  |  | VP | -1.21 | 0.51 |
|  |  | CBCL | 1.5 | 0.17 |
|  |  | BRIEF | 1.34 | 0.18 |
|  |  | ABAS | -1.16 | 0.60 |
|  |  | NIHTB Flanker | -1.51 | 0.09 |
|  |  | NIHTB Card Sort | -1.39 | 0.27 |
|  | P2-N2 | NVIQ | 0.53 | 0.49 |
|  |  | Language | 1.79 | 0.01* |
|  |  | VMI | -0.69 | 0.11 |
|  |  | VP | -1.06 | 0.45 |
|  |  | CBCL | 1.13 | 0.15 |
|  |  | BRIEF | 0.73 | 0.26 |
|  |  | ABAS | -1.2 | 0.50 |
|  |  | NIHTB Flanker | -0.99 | 0.08 |
|  |  | NIHTB Card Sort | -0.72 | 0.37 |
|  | N2 | NVIQ | -0.75 | 0.64 |
|  |  | Language | -3.33 | 0.02* |
|  |  | VMI | 2.84 | 0.01* |
|  |  | VP | 3.29 | 0.44 |
|  |  | CBCL | -1.35 | 0.20 |
|  |  | BRIEF | -0.91 | 0.53 |
|  |  | ABAS | 5.07 | 0.34 |
|  |  | NIHTB Flanker | 1.79 | 0.16 |
|  |  | NIHTB Card Sort | -3.12 | 0.47 |
| RSTS | N1 | NVIQ | -2.51 | 0.42 |
|  |  | Language | -4.61 | 0.12 |
|  |  | VMI | -0.18 | 0.95 |
|  |  | VP | -5.65 | 0.11 |
|  |  | CBCL | -1.3 | 0.49 |
|  |  | BRIEF | -0.34 | 0.87 |
|  |  | ABAS | 1.61 | 0.60 |
|  |  | NIHTB Flanker | -7.35 | 0.08 |
|  |  | NIHTB Card Sort | -2.24 | 0.54 |
|  | N1-P2 | NVIQ | 2.07 | 0.24 |
|  |  | Language | 2.28 | 0.10 |
|  |  | VMI | 1.84 | 0.26 |
|  |  | VP | 3.96 | 0.11 |
|  |  | CBCL | 1.1 | 0.29 |
|  |  | BRIEF | 1.54 | 0.24 |
|  |  | ABAS | -1.5 | 0.38 |
|  |  | NIHTB Flanker | 3.35 | 0.05 |
|  |  | NIHTB Card Sort | 1.28 | 0.40 |
|  | P2 | NVIQ | 2.9 | 0.29 |
|  |  | Language | 3.27 | 0.13 |
|  |  | VMI | 4.61 | 0.06 |
|  |  | VP | 4.24 | 0.30 |
|  |  | CBCL | 1.59 | 0.32 |
|  |  | BRIEF | 3.04 | 0.08 |
|  |  | ABAS | -2.3 | 0.38 |
|  |  | NIHTB Flanker | 4.6 | 0.06 |
|  |  | NIHTB Card Sort | 2.16 | 0.30 |
|  | P2-N2 | NVIQ | 1.29 | 0.43 |
|  |  | Language | 1.17 | 0.27 |
|  |  | VMI | 1.79 | 0.22 |
|  |  | VP | 1.47 | 0.52 |
|  |  | CBCL | 0.71 | 0.41 |
|  |  | BRIEF | 1.85 | 0.07 |
|  |  | ABAS | -1.22 | 0.37 |
|  |  | NIHTB Flanker | 4.19 | 0.07 |
|  |  | NIHTB Card Sort | 1.93 | 0.33 |
|  | N2 | NVIQ | -1.52 | 0.63 |
|  |  | Language | -1.78 | 0.48 |
|  |  | VMI | -1.44 | 0.61 |
|  |  | VP | -0.71 | 0.87 |
|  |  | CBCL | -0.94 | 0.56 |
|  |  | BRIEF | -3.34 | 0.09 |
|  |  | ABAS | 2.1 | 0.42 |
|  |  | NIHTB Flanker | -6.1 | 0.42 |
|  |  | NIHTB Card Sort | -3.71 | 0.54 |
| KS + RSTS | N1 | NVIQ | -4.03 | 0.08 |
|  |  | Language | -2.14 | 0.19 |
|  |  | VMI | -2.17 | 0.25 |
|  |  | VP | -4.85 | 0.09 |
|  |  | CBCL | -2.12 | 0.06 |
|  |  | BRIEF | -1.03 | 0.25 |
|  |  | ABAS | -0.55 | 0.84 |
|  |  | NIHTB Flanker | -4.13 | 0.02* |
|  |  | NIHTB Card Sort | 2.22 | 0.16 |
|  | N1-P2 | NVIQ | 1.52 | 0.13 |
|  |  | Language | 1.32 | 0.04* |
|  |  | VMI | 0.9 | 0.22 |
|  |  | VP | 1.19 | 0.17 |
|  |  | CBCL | 0.97 | 0.07 |
|  |  | BRIEF | 0.97 | 0.05* |
|  |  | ABAS | -0.05 | 0.95 |
|  |  | NIHTB Flanker | 1.65 | 0.01* |
|  |  | NIHTB Card Sort | -1.31 | 0.14 |
|  | P2 | NVIQ | 1.99 | 0.22 |
|  |  | Language | 2.73 | 0.01* |
|  |  | VMI | 1.46 | 0.21 |
|  |  | VP | 1.38 | 0.35 |
|  |  | CBCL | 1.49 | 0.08 |
|  |  | BRIEF | 1.86 | 0.03* |
|  |  | ABAS | -0.17 | 0.88 |
|  |  | NIHTB Flanker | 2.5 | 0.01* |
|  |  | NIHTB Card Sort | 1.81 | 0.17 |
|  | P2-N2 | NVIQ | 0.82 | 0.25 |
|  |  | Language | 1.66 | 0.004* |
|  |  | VMI | -0.42 | 0.17 |
|  |  | VP | -0.26 | 0.49 |
|  |  | CBCL | 1.11 | 0.09 |
|  |  | BRIEF | 1.11 | 0.02* |
|  |  | ABAS | -0.48 | 0.72 |
|  |  | NIHTB Flanker | 1.68 | 0.03* |
|  |  | NIHTB Card Sort | 1.22 | 0.23 |
|  | N2 | NVIQ | -1.24 | 0.28 |
|  |  | Language | -2.83 | 0.02* |
|  |  | VMI | 1.8 | 0.02* |
|  |  | VP | 0.79 | 0.41 |
|  |  | CBCL | -1.15 | 0.18 |
|  |  | BRIEF | -1.77 | 0.06 |
|  |  | ABAS | 4.29 | 0.35 |
|  |  | NIHTB Flanker | 1.79 | 0.16 |
|  |  | NIHTB Card Sort | -2.68 | 0.39 |
